# Towards a successful translocation of captive slow lorises (*Nycticebus* spp.) in Borneo: a review and recommendations

**DOI:** 10.1101/078535

**Authors:** C. M. G. van der Sandt

**Affiliations:** Orangutan Project Sdn Bhd., Kuching, Sarawak, Borneo.

**Keywords:** reintroduction, prosimian, rehabilitation, conservation

## Abstract

Wildlife in Southeast Asia is under threat mainly due to habitat loss and the illegal animal trade. Wild animals are rescued by wildlife centres and the slow loris (*Nycticebus* spp.) is one of them. After rehabilitation slow lorises are released into the wild and on average 26% is observed alive after at least ten weeks.

The IUCN has established conditions under which captive wild animals can be translocated into their natural habitat. This review paper aims to give an overview of what has been published on release programs of captive slow lorises in order to improve translocations in Sarawak, Malaysian Borneo. Five documented studies on translocation of slow lorises are summarised. I concentrated on: (1) species of captive slow loris, (2) health check, (3) pre-release habituation, (4) soft or hard release, (5) pre and post-release behavioural observations, (6) assessment of the release area: predators, habitat and protection. The recommendations for future releases are: (1) Study which slow loris species are rehabilitated in Bornean wildlife centres. (2) Study the behaviour of captive slow lorises. (3) Assess the slow loris species in the release area. (4) Study the behaviour and habitat use of the wild population. (5) Assess what predators are present in the release area.

## INTRODUCTION

In Southeast Asia large numbers of wild animals are rehabilitated in wildlife and rescue centres (Collins *et al*., 2008; Moore *et al*., 2014). Wildlife centres care for a.o. orangutans, macaques, slow lorises, wild cats, birds, snakes, turtles, binturongs and sun bears (Isler & Thorpe, 2003; International Animal Rescue, 2006-2016; Biddle, 2015; pers. obs. CvS, 2009-2015). In the wild, Bornean animal species are threatened in different ways; animals are captured to be sold and kept as pets or used in traditional medicines (Shepherd *et al*., 2004) and rainforest is converted to farmland and oil palm plantations (Nekaris & Starr, 2015). Wild animals cannot survive in these new habitats. They are killed or are caught and sold (Shepherd *et al*., 2004). It is illegal to catch and keep many of the wild Bornean animal species and captive animals are confiscated by the government and transferred to wildlife centres. Furthermore, the centres are also offered captive wild animals, which were purchased in animal markets by local people or tourists and are no longer wanted (pers. obs. CvS, 2015). The aim of wildlife centres is to rehabilitate the animals and, if possible, release them back into the wild (Biddle, 2015; Nekaris & Starr, 2015). The International Union for Conservation of Nature (IUCN) has established conditions under which captive wild animals can be released or translocated into their natural habitat (IUCN, 2002; IUCN/SSC, 2013). According to the IUCN, translocation is “*the human-mediated movement of living organisms from one area, with release in another*”.

### The slow loris

The slow loris (*Nycticebus* spp.) is a small nocturnal strepsirrhine primate from Southeast Asia. The average weight of an adult slow loris is between 265 and 1150 grams, depending on the species (Nekaris, 2014). They have a round head and large forward-facing eyes. The arms and legs are adapted to an arboreal life. Slow lorises are the only venomous primates; they have a brachial gland that produces a secretion which is toxic when licked by the slow loris and mixed with saliva. The bite is poisonous to other animals (Ligabue-Braun *et al*., 2012; Rode-Margono & Nekaris, 2015). Despite their protected status, slow lorises are widely caught and sold in markets throughout Southeast Asia. They are very popular as pets and are used in traditional medicines (Nekaris & Starr, 2015).

Nine species of slow loris have been identified (Nekaris *et al*., 2014): The Bengal slow loris (*N. bengalensis* Lacépède), Sunda or greater slow loris (*N. coucang* Boddaert), Hiller’s slow loris (*N. hilleri* Stone & Rehn), Javan slow loris (*N. javanicus* É. Geoffroy) and pygmy slow loris (*N. pygmaeus* Bonhote). Recently four Bornean species of slow loris have been recognized: Sody’s slow loris (*N. bancanus* Lyon), Bornean slow loris *(N. borneanus* Lyon), Kayan River slow loris *(N. kayan* Munds, Nekaris & Ford) and Philippine slow loris (*N. menagensis* Lydekker) (Munds *et al*., 2013). Different species of slow loris are kept in wildlife centres. It is hard to keep slow lorises in captivity for a long period, due to incurred stress, their nocturnal lifestyle and their diet (Beckerson, 2015; Nekaris & Starr, 2015). If released into their natural habitat, many of the slow lorises do not survive for a very long time (Moore *et al*., 2014; Biddle, 2015; Nekaris & Starr, 2015).

### Slow lorises in Matang Wildlife Centre, Malaysian Borneo

Since 2009 there have been 29 documented arrivals of slow lorises at Matang Wildlife Centre. Seven of these arrivals were released within two to three days, and four within two to four months (reference date April 2017) (Beckerson, 2016; Browning, 2017). Matang is part of Kubah National Park, Sarawak, Malaysia. Kubah, 15 km northwest of Kuching, is 22 sq. km in size. The bulk of the park is a sandstone plateau at an elevation between 150 and 450 m asl and it has five main vegetation types: riverine forest, lowland mixed dipterocarp forest, kerangas (heath) forest, submontane forest and secondary forest.

**Fig. 1.**
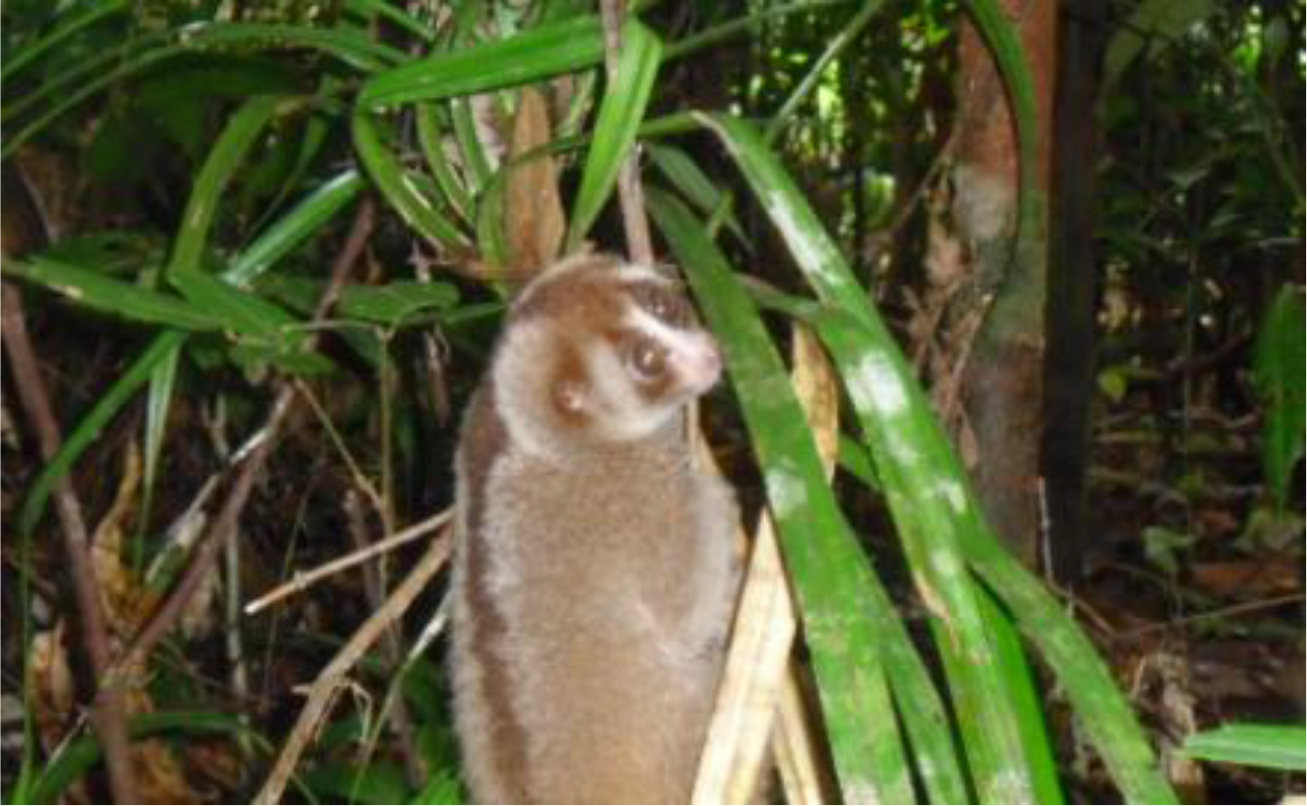
Slow loris in Matang Wildlife Centre (Aug 2009) Photo: Carola Voigt

In this paper I aim to give an overview of what has been published (reference date August 2016) on rehabilitation and release programs of captive slow lorises, in order to increase the number of future successful translocations in Sarawak, Malaysian Borneo. The four main questions of this review paper are:

1. *Which slow loris species have been studied?* Nine slow loris species have been described (Nekaris *et al*., 2014). Although there are some overlaps, these species are found in different areas and habitats and have different feeding requirements and behaviour. It is important to know the species of captive slow loris, in order to supply the animals with the most suitable food during rehabilitation and to assess the area and habitat where the lorises will be released.
2. *What pre- and post-conditions for release have to be met for the best possible success for survival after release?* As the main objective of this paper is to identify the preconditions to increase the number of future successful translocations, it is important to know which preconditions researchers have used, in relation to the percentage of surviving animals (for any length of time) in their studies. This argument leads to the following questions:
3. *What percentage of slow lorises did survive after release?*
4. *What recommendations did the researchers make for future research?* The release of slow lorises faces many challenges. The recommendations of previous studies might help to improve the results of future translocations.

## METHODS

Literature was collected between February and May 2016. Online data bases were searched for articles, books and doctoral theses in English: Conservation database for lorises (*Loris, Nycticebus*) and pottos (*Arctocebus, Perodicticus*), prosimian primates; Elsevier Science Direct; Google Scholar. A web search was carried out with the following keywords: slow loris, captive slow loris, conservation, rehabilitation, translocation, reintroduction, primates. The snowball method was used. L. Biddle and N. Beckerson, respectively founder and manager of the Orangutan Project, based in Matang Wildlife Centre, Sarawak, Malaysia, answered questions for clarification of results found, both in person and via email. This paper is a review of documented research on rehabilitation and release of captive wild-born slow lorises, in accordance with the IUCN Guidelines. The IUCN issued two guidelines on translocations and reintroductions: Guidelines for nonhuman primate re-introductions (2002); Guidelines for Reintroductions and Other Conservation Translocations (2013). The regulations in these documents that are within the scope of this paper include: (1) Basic biology of the species, biotic and abiotic habitat needs, interspecific relationships; (2) Quarantine before release; (3) Welfare of the captive animal to avoid stress during rehabilitation, handling and transport; (4) Health and behaviour, both of the captive animal that is going to be released and of the wild population in the release area; (5) Assessment of suitable release habitat, niches of the translocated species; (6) The season of release; (7) Pre- and post-release monitoring of the animals and the release site.

The use of the following terms in this paper is based on the IUCN Guidelines and Moore *et al*. (2014). <u>Release</u>: a general term to indicate the process of release into the wild, rehabilitation and monitoring excluded. <u>Translocation</u>: the human-mediated movement of living organisms from one area, with release in another. <u>Conservation translocation</u>: a translocation to reinforce an existing population of conspecifics. <u>Reintroduction</u>: the reintroduction of a species into an area which was once part of its range, but from which it has been extirpated or become extinct.

The last decade has seen an increasing number of slow lorises released by wildlife centres. Many of the slow lorises have simply been released into any suitable habitat in the surroundings of the centre without a thorough health check, species study or knowledge of their ability to survive in the wild (Streicher, 2004; Moore *et al*., 2014; Nekaris & Starr, 2015). These often unpublished incidences of release are not included in this paper.

## RESULTS

The first question was which slow loris species have been studied and what was the number of translocated animals. Table 1 shows five documented studies of three slow loris species: the Javan slow loris (*Nycticebus javanicus*), pygmy slow loris (*N. pygmaeus*) and Sunda or greater slow loris (*N. coucang*). The number of translocated slow lorises in the five studies ranges from 5 to 18 specimens.

**Table 1.**
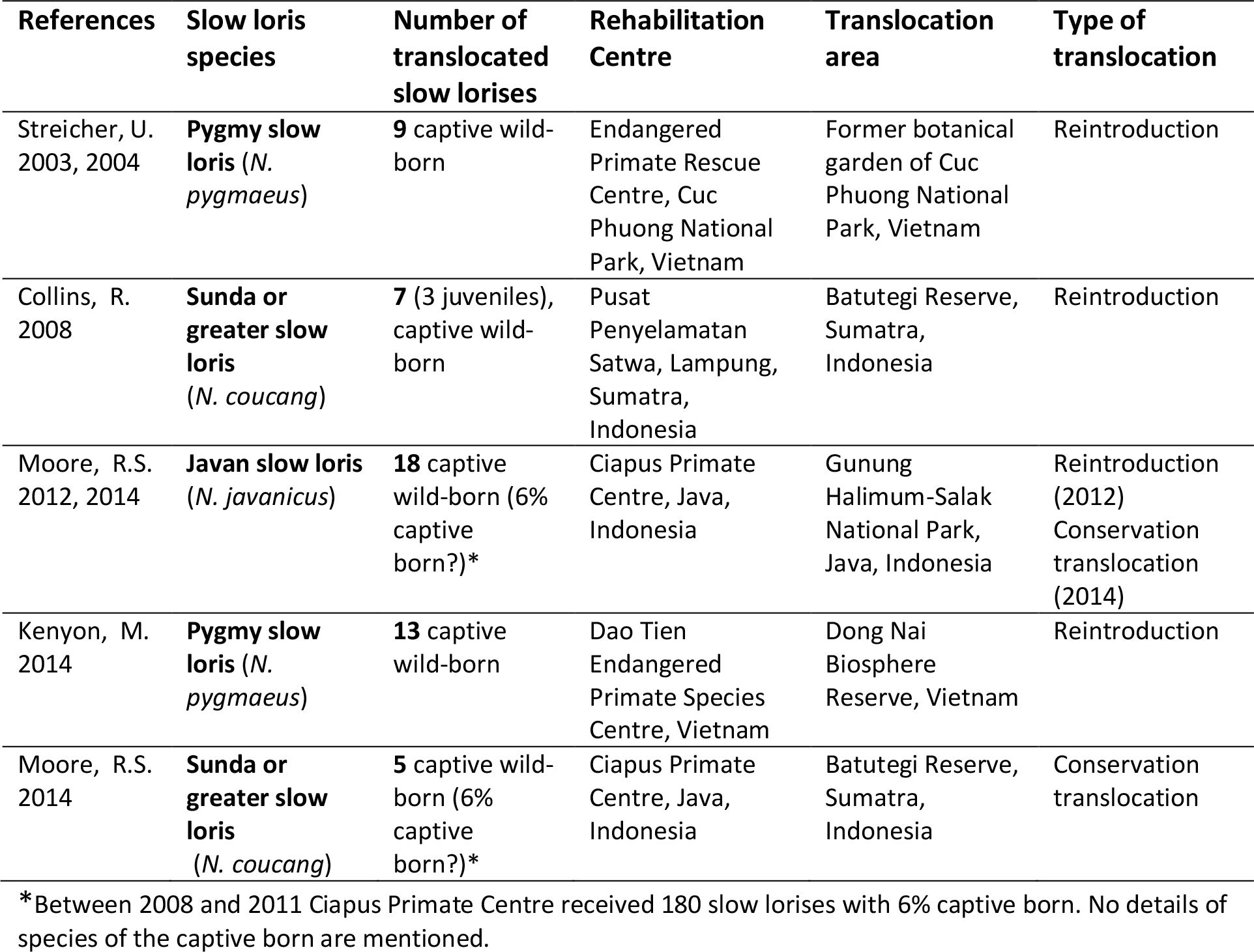
Documented translocation studies of captive wild-born slow lorises (*Nycticebus* spp.) in Indonesia and Vietnam.

Moore (2014) states that over 85% of the confiscated animals in his study were unsuitable for release. In many captive lorises, the teeth are cut or removed before confiscation to prevent the lorises from biting during handling in animal markets. All slow loris species are obligate gum eaters (Nekaris, 2014). Gum is acquired by the gouging of tree bark. Slow lorises without teeth will never be able to gouge for gum, so are unsuitable for release. Stress and very poor living conditions before entering the rescue centres are also causes of permanent physical and behavioural damage. This results in a low number of animals suitable for translocation. The authors describe their releases as reintroductions or conservation translocations. The second and third questions concerned the pre- and post-conditions for release and the percentage of surviving slow lorises after release.

**Table 2.**
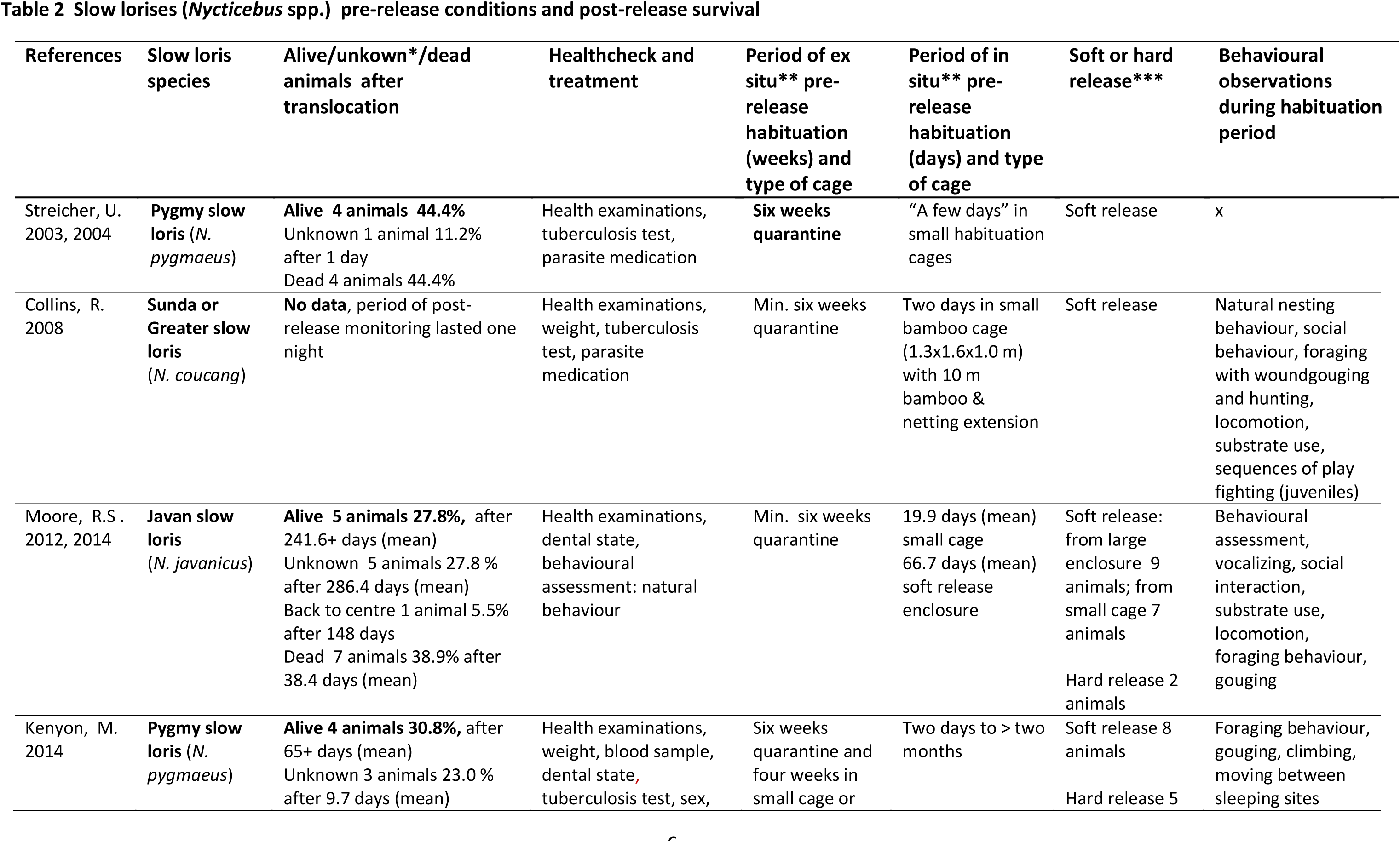
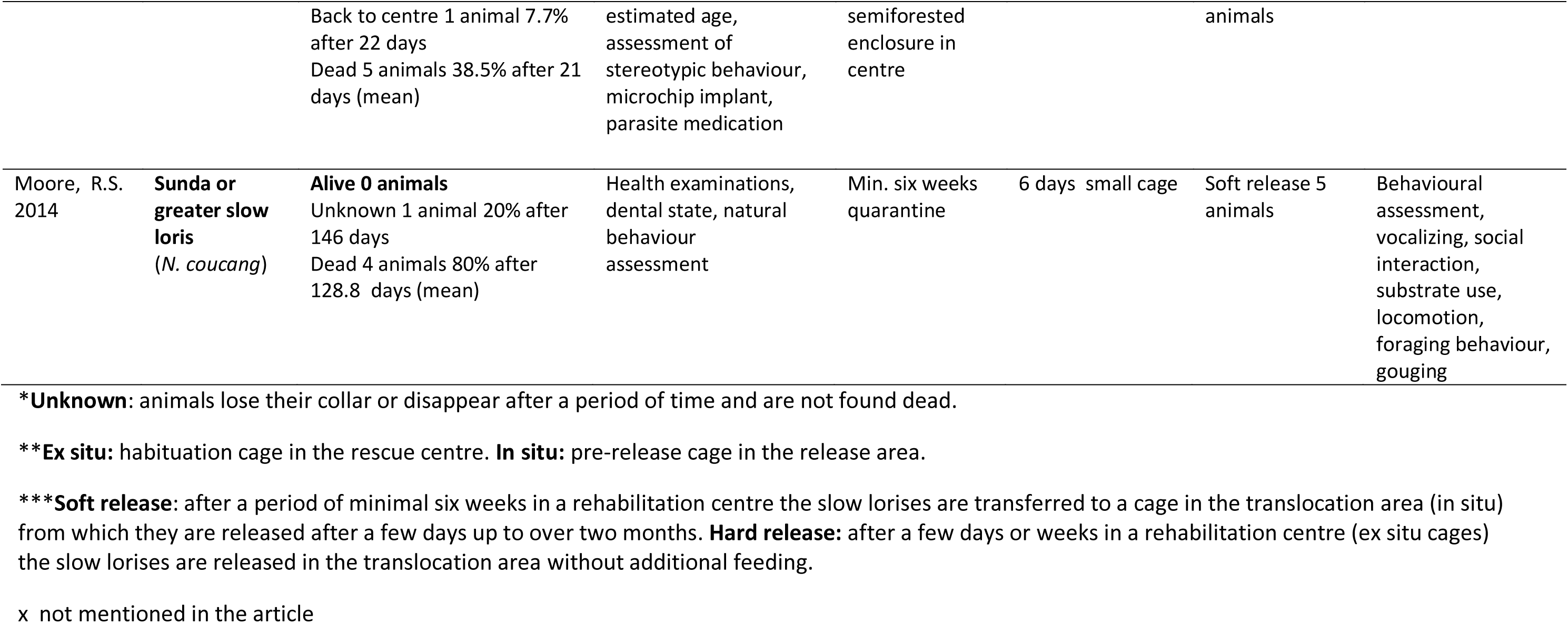
Slow lorises (*Nycticebus* spp.) pre-release conditions and post-release survival summarizes the pre-release preconditions during rehabilitation and the percentage of animals that survived after release.

| References | Slow loris species | Alive/unknown*/dead animals after translocation | Healthcheck and treatment | Period of ex situ** pre-release habituation (weeks) and type of cage | Period of in situ** pre-release habituation (days) and type of cage | Soft or hard release*** | Behavioural observations during habituation period |
| --- | --- | --- | --- | --- | --- | --- | --- |
| Streicher, U. 2003, 2004 | **Pygmy slow loris** (*N. pygmaeus*) | **Alive 4 animals 44.4%**<br>Unknown 1 animal 11.2% after 1 day<br>Dead 4 animals 44.4% | Health examinations, tuberculosis test, parasite medication | **Six weeks quarantine** | "A few days" in small habituation cages | Soft release | x |
| Collins, R. 2008 | **Sunda or Greater slow loris** (*N. coucang*) | **No data**, period of post-release monitoring lasted one night | Health examinations, weight, tuberculosis test, parasite medication | Min. six weeks quarantine | Two days in small bamboo cage (1.3x1.6x1.0 m) with 10 m bamboo & netting extension | Soft release | Natural nesting behaviour, social behaviour, foraging with woundgouging and hunting, locomotion, substrate use, sequences of play fighting (juveniles) |
| Moore, R.S. 2012, 2014 | **Javan slow loris** (*N. javanicus*) | **Alive 5 animals 27.8%**, after 241.6+ days (mean)<br>Unknown 5 animals 27.8 % after 286.4 days (mean)<br>Back to centre 1 animal 5.5% after 148 days<br>Dead 7 animals 38.9% after 38.4 days (mean) | Health examinations, dental state, behavioural assessment: natural behaviour | Min. six weeks quarantine | 19.9 days (mean) small cage<br>66.7 days (mean) soft release enclosure | Soft release: from large enclosure 9 animals; from small cage 7 animals<br>Hard release 2 animals | Behavioural assessment, vocalizing, social interaction, substrate use, locomotion, foraging behaviour, gouging |
| Kenyon, M. 2014 | **Pygmy slow loris** (*N. pygmaeus*) | **Alive 4 animals 30.8%**, after 65+ days (mean)<br>Unknown 3 animals 23.0 % after 9.7 days (mean) | Health examinations, weight, blood sample, dental state, tuberculosis test, sex, | Six weeks quarantine and four weeks in small cage or | Two days to > two months | Soft release 8 animals<br>Hard release 5 | Foraging behaviour, gouging, climbing, moving between sleeping sites |
|  |  | Back to centre 1 animal 7.7% after 22 days<br>Dead 5 animals 38.5% after 21 days (mean) | estimated age, assessment of stereotypic behaviour, microchip implant, parasite medication | semiforested enclosure in centre |  | animals |  |
| Moore, R.S. 2014 | **Sunda or greater slow loris**<br>(*N. coucang*) | **Alive 0 animals**<br>Unknown 1 animal 20% after 146 days<br>Dead 4 animals 80% after 128.8 days (mean) | Health examinations, dental state, natural behaviour assessment | Min. six weeks quarantine | 6 days small cage | Soft release 5 animals | Behavioural assessment, vocalizing, social interaction, substrate use, locomotion, foraging behaviour, gouging |

\***Unknown:** animals lose their collar or disappear after a period of time and are not found dead.
\*\***Ex situ:** habituation cage in the rescue centre. **In situ:** pre-release cage in the release area.
\*\*\***Soft release:** after a period of minimal six weeks in a rehabilitation centre the slow lorises are transferred to a cage in the translocation area (in situ) from which they are released after a few days up to over two months. **Hard release:** after a few days or weeks in a rehabilitation centre (ex situ cages) the slow lorises are released in the translocation area without additional feeding.
x not mentioned in the article

Survival rates are between 0 and 44.4%. In Kenyon’s study (2014) four animals survived. They all had a soft release. When hard released the fate of the animals is unknown or the animals died. The pre-release preconditions matched most of the guidelines of the IUCN (2002, 2013): (1) A health check of the captive animals was done in all studies. As stated above, the dental state of slow lorises is important. In one study no check of dental state or referral to ability to gouge is mentioned. However, after release tree gouging was observed and the thesis stresses the importance of the toothcomb in the lower jaw (Streicher, 2004); (2) Quarantine periods of a minimum of six weeks is mentioned in all studies; (3) Welfare of the captive animals to avoid stress was guaranteed by pre-release habituation. There are differences between the studies: from ex situ habituation in small cages and a subsequent hard release, to ex situ and in situ habituation in semi forested enclosures (soft release); (4) Behavioural observations during the pre-release habituation period is mentioned in four studies. The success of a translocation project can be measured by the number of animals surviving in a given period.

Figure 2 summarises the percentage of surviving slow lorises during the observation period in three studies (Kenyon, 2014; Moore, 2012; Moore *et al*., 2014). In two studies no detailed information on this topic is available or applicable (Streicher, 2003; Collins, 2008).

**Fig. 2.**
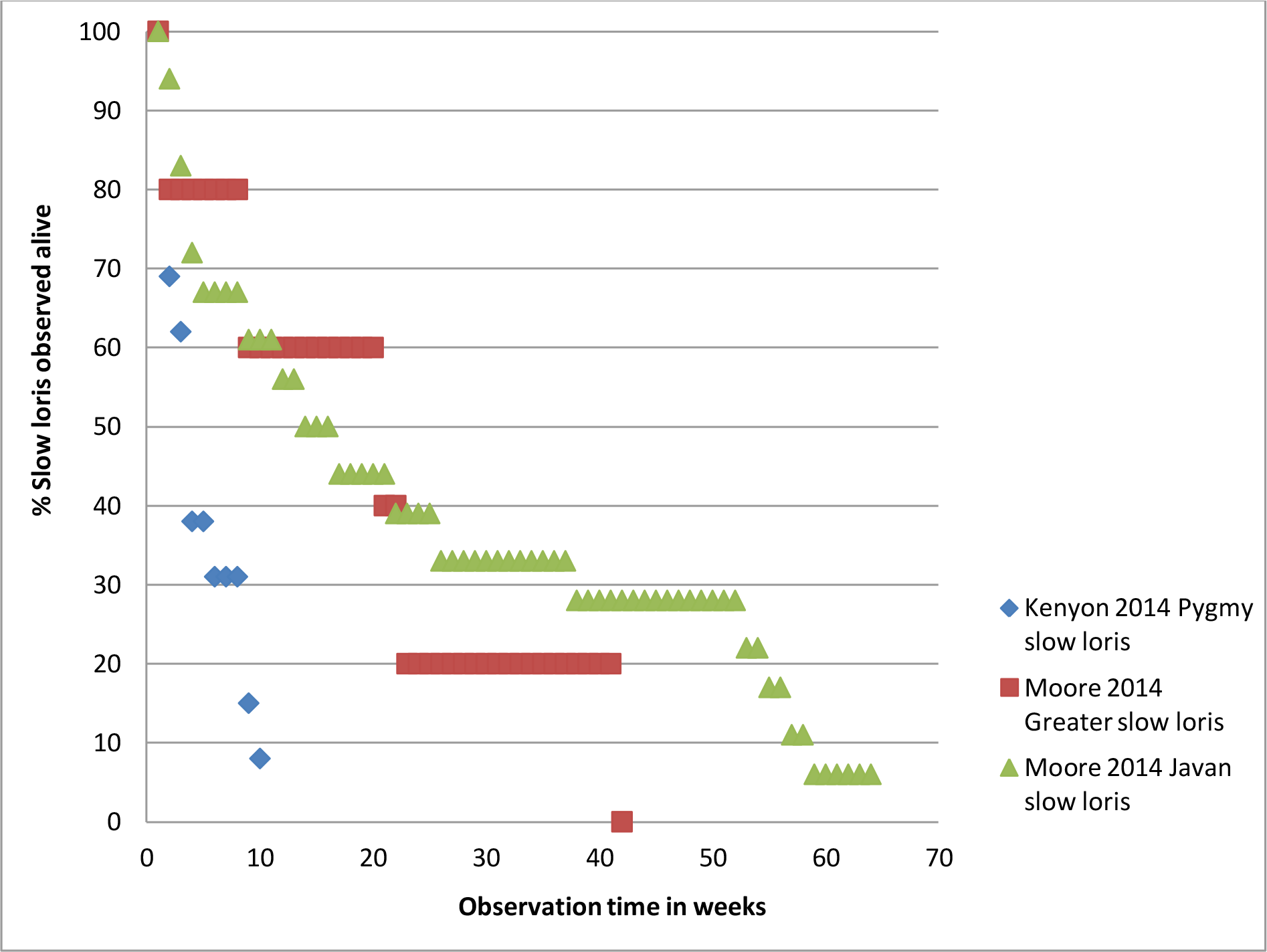
Percentage slow loris survival in observation period.

**Table 3.** Pre-release preconditions of the translocation area and post-release survival of slow lorises shows an overview of the pre-release habitat preconditions and the post-release survival of slow lorises.

| References | Slow loris species | Alive/unknown*/dead animals after translocation |  |  | Existing population of slow lorises in translocation area | Assessment of predators in translocation area | Habitat/vegetation assessment | Protection translocation area |
| --- | --- | --- | --- | --- | --- | --- | --- | --- |
| Streicher, U. 2003, 2004 | **Pygmy slow loris**<br>(*N. pygmaeus*) | **Alive** | **4 animals** | **44.4%** | One resident pygmy slow loris in 1999 | x | Dense vegetation | National park |
|  |  | Unknown | 1 animal | 11.2% |  |  |  |  |
|  |  | Dead | 4 animals | 44.4% |  |  |  |  |
| Collins, R. 2008 | **Sunda or greater slow loris**<br>(*N. coucang*) | **No data**, period of post-release monitoring lasted one night |  |  | “Low abundance of lorises” in 2006 | x | x | Limited human impact, high level of protection by Forest Department |
| Moore, R.S. 2012, 2014 | **Javan slow loris**<br>(*N. javanicus*) | **Alive** | **5 animals** | **27.8%** | Assessment of slow lorises in 2007, 2011 | x | Habitat assessment | National park |
|  |  | Unknown | 5 animals | 27.8% |  |  |  |  |
|  |  | Dead | 7 animals | 38.9% |  |  |  |  |
|  |  | Back to centre | 1 animal | 5.5% |  |  |  |  |
| Kenyon, M. 2014 | **Pygmy slow loris**<br>(*N. pygmaeus*) | **Alive** | **4 animals** | **30.8%** | Assessment of pygmy lorises in one release site (Cat Tien National Park), not in Vinh Cuu Biosphere Reserve | Estimation of predator density | Forest connectivity, percentage of ground cover, tree occupation | National park |
|  |  | Unknown | 3 animals | 23.0% |  |  |  |  |
|  |  | Dead | 5 animals | 38.5% |  |  |  |  |
|  |  | Back to centre | 1 animal | 7.7% |  |  |  |  |
| Moore, R.S. 2014 | **Sunda or greater slow loris**<br>(*N. coucang*) | **Alive** | **0 animals** |  | Assessment of slow lorises | x | Habitat assessment | National reserve |
|  |  | Unknown | 1 animal | 20% |  |  |  |  |
|  |  | Dead | 4 animals | 80% |  |  |  |  |

\*Unknown: animals may lose their collar or disappear after a period of time and are not found dead.
x not mentioned in the article

(5) An assessment of an existing population of slow lorises in the translocation area is done in all but one translocation area where permission was withheld by the local authorities (Kenyon *et al*., 2014); (6) The assessment of predators in the translocation area is listed separately in the present paper as it is of significant importance (pers. obs. CvS, 2015). Four studies do not mention a pre-release assessment of predators, although they were present in the release area. In one study the attack of slow lorises by snakes and probably a raptor in the release area is observed (Moore *et al*., 2014). In Streicher’s study two lorises were killed by predators, one of which was a marbled cat (Streicher & Nadler, 2003). (7) An assessment of the vegetation is carried out in four studies and all translocation areas are within national parks, so a reasonable level of protection must be assumed. Table 4 is a summary of the post-release monitoring of slow lorises related to the surviving animals.

**Table 4.**
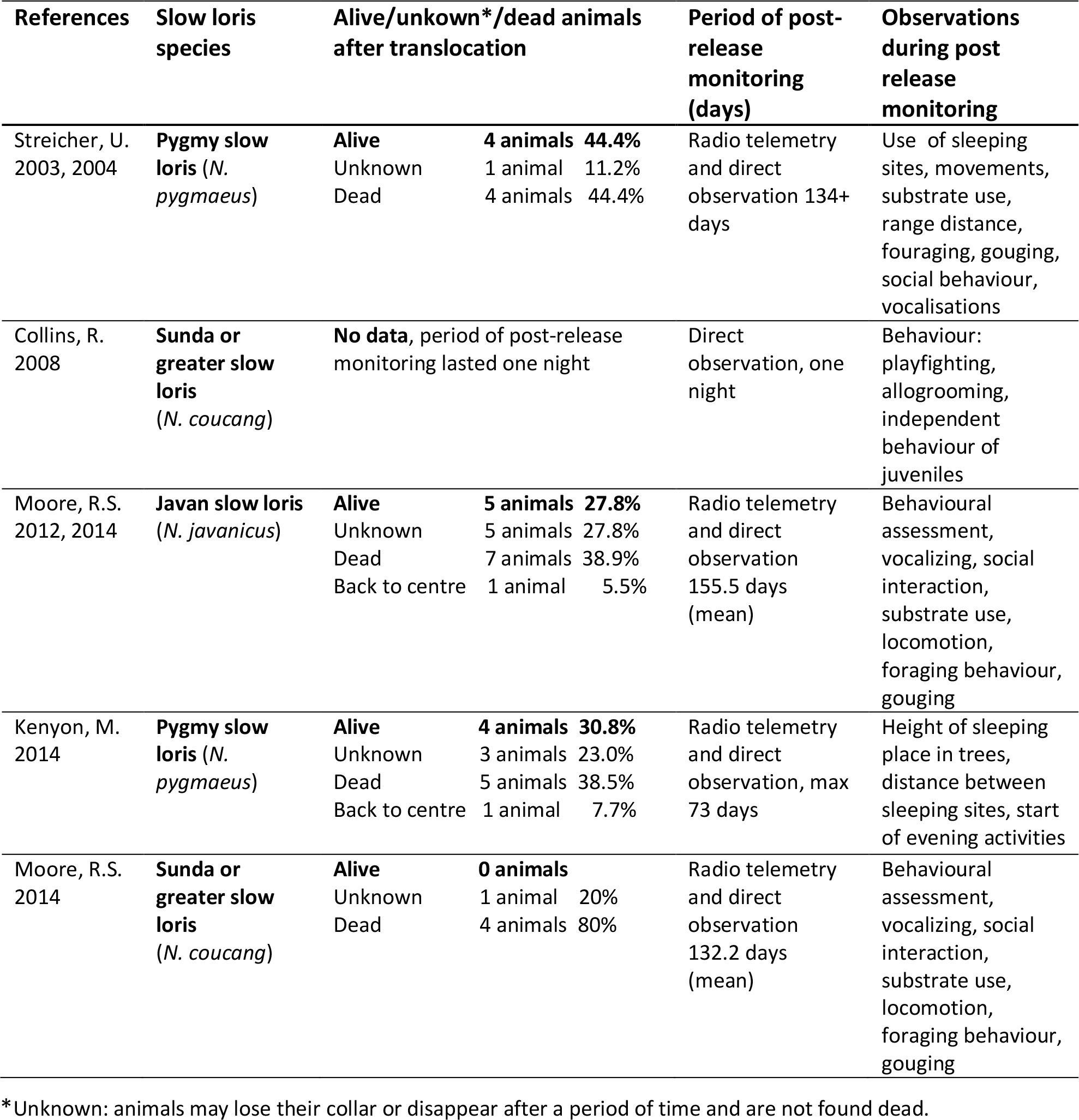
Post-release preconditions and post-release survival of slow lorises: the animals.

(8) In one study post-release monitoring lasted only one night, due to local customs regarding entering the forest at night and communication on ecological issues with local workers (Collins *et al*., 2008); (9) In all studies post-release behaviour of the slow lorises is studied. Four studies used direct observation and radio telemetry. In one of the translocations no radio telemetry was applied (Collins *et al*., 2008); (10) A health check of resident slow lorises is not applied in any of the studies. The IUCN guidelines of 2013 however, stress the importance of an assessment of the health of wild populations. Regarding the final research question on recommendations, Table 5 summarizes the recommendations for future research as mentioned by the cited authors. Also included are suggestions presented in an overview paper on conservation and ecology of slow lorises by Nekaris and Starr (2015).

**Table 5.**
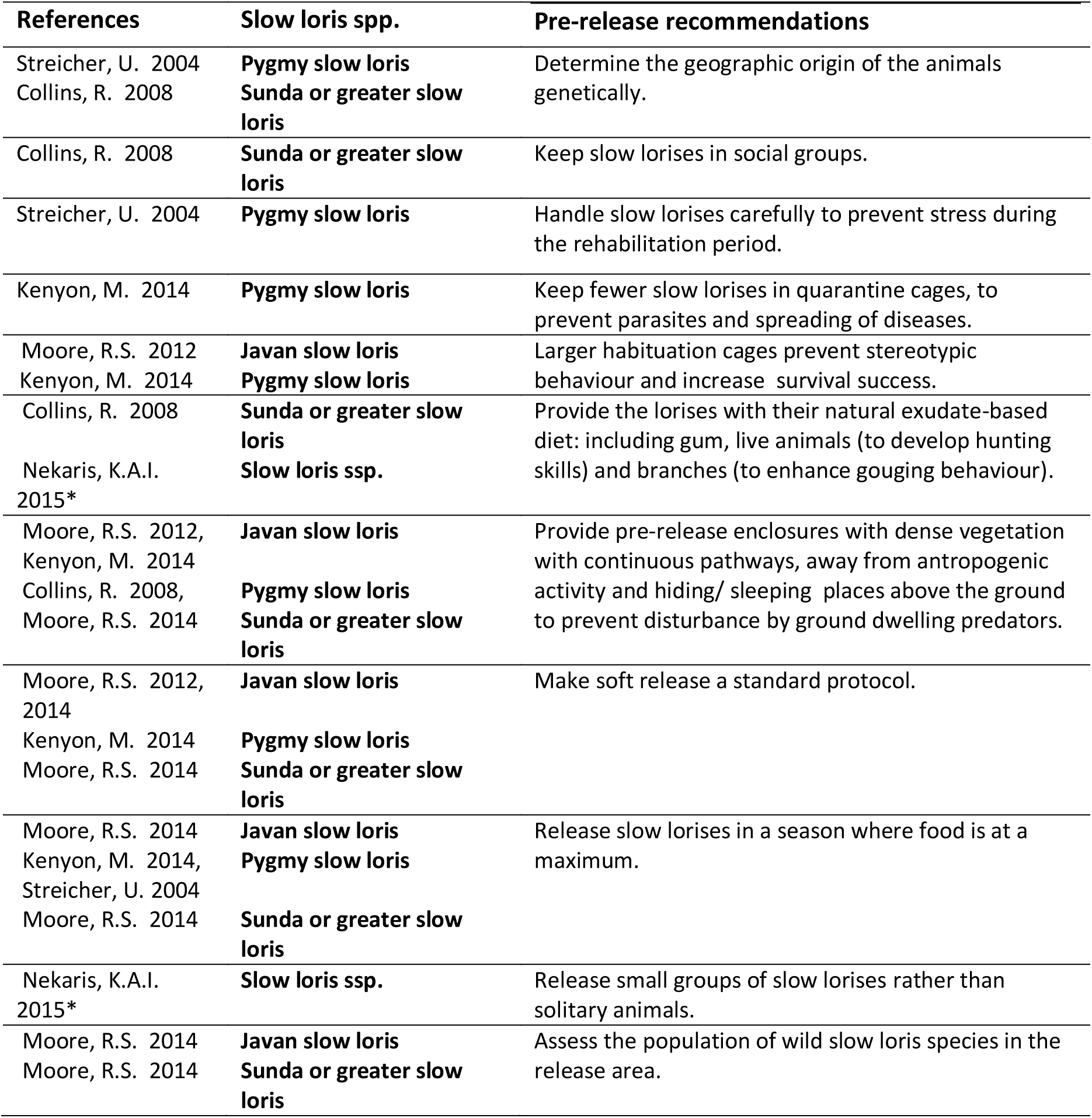
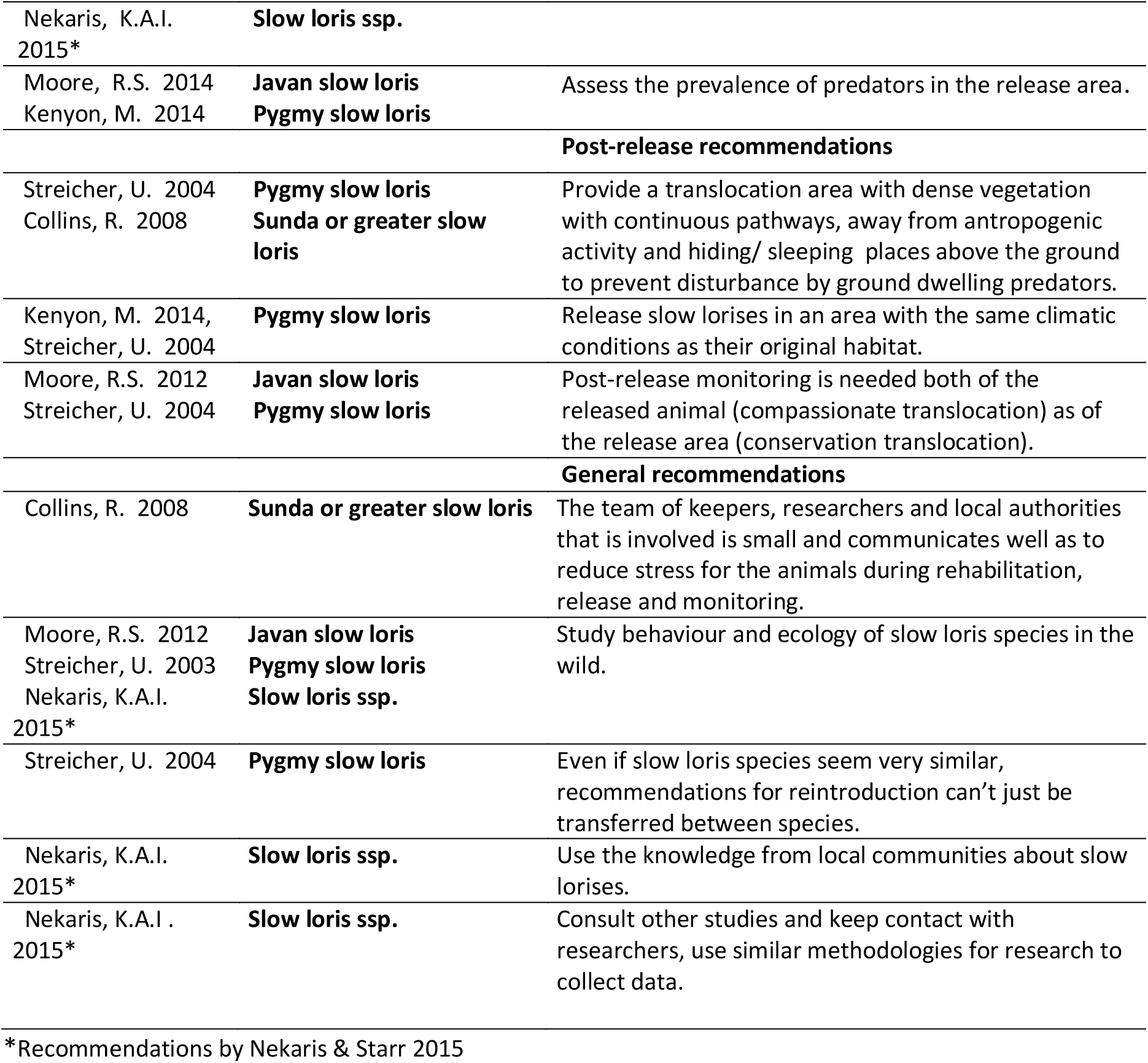
Recommendations for future research on rehabilitation and translocation of slow lorises.

Recommendations mentioned in at least three references were: (1) Provide the pre-release enclosures with dense vegetation with continuous pathways, away from anthropogenic activity and hiding/ sleeping places above the ground to prevent disturbance by ground dwelling predators. (2) Make soft release a standard protocol. (3) Release slow lorises in a season where food is at a maximum. (4) Study the behaviour and ecology of slow loris species in the wild.

## DISCUSSION

This review paper aimed to make recommendations for rehabilitation and translocation of the slow loris species (*Nycticebus* ssp.) in Malaysian Borneo. A limitation could be that the slow loris species of the studies in this review are not native to Malaysian Borneo. However, the different species have a lot in common. The findings in the present paper could therefore still be valuable to the Bornean situation. Another limitation was that all the analyzed studies involve only small numbers of released animals. However, it was still useful to compare these studies to discover possible trends. Figure 2 compares the success of three translocation studies. The reintroduction of the Javan slow loris is most successful (Moore *et al*., 2014). Moore (2014) mentions a slight improvement of post-release survival of the Javan slow loris in his paper as well. The main difference between the Moore and Kenyon studies is the use of a large habituation cage in 50% of the releases of Javan slow lorises by Moore. A point of attention is that in wild populations a certain percentage of animals die, due to predation, hypothermia, lack of food, hunting etcetera. Goodman (1993) estimated death rates due to predation alone in *Microcebus* populations to be 25%. The number of animals in wild slow loris populations that dies due to predation might also be high (Goodman *et al*., 1993). Assessment of predators in the release area is important because in the studies of Moore (2014) and Streicher (2003) a number of released animals died due to predation. Most predation of slow loris occurs when they are forced to move between trees on the forest floor. Ground cover in a large habituation enclosure and release area, and dense vegetation with continuous pathways well above the ground are therefore of major importance. Radio telemetry supplemented by direct observation seems to also be important. The animals spend time in thick vegetation, making direct observation impossible. Regarding the recommendations that were found, it is important to understand whether and how local workers want to participate in a research program.

In addition to the recommendations for rehabilitation and release, mentioned in the IUCN guidelines of 2002 and 2013, the following recommendations, some of which are also mentioned by the IUCN, are of interest to wildlife centres in Malaysian Borneo: (1) Study which slow loris species are rehabilitated in Bornean wildlife centres; (2) Study the behavioural characteristics of the captive slow lorises in Bornean wildlife centres; (3) Assess the wild slow loris species living in the release area; (4) Study the behaviour and habitat use of the wild population; (5) Assess what predators are present in the release area.

It might also be valuable to assess in a computer simulation what percentage of a natural population of slow lorises dies due to different factors such as old age, disease, weather conditions, predation and food conditions in their habitat.

Following these recommendations will hopefully lead to a better conservation of these beautiful species.

## Acknowledgements

This paper benefited enormously from the contributions of the following people: N. Beckerson and L. Biddle provided valuable information. V. P. J. Borm and M. Hopman-Rock greatly improved the quality of this manuscript. B.G. de Vries commented on the manuscript. I.E.E. Scheper provided valuable literature. Thanks to C. Voigt who shared the photo of *Nycticebus* sp. Additional thanks to one anonymous reviewer.

